# Epigenetic regulation of frog innate immunity: Discovery of frog microRNAs associated with antiviral responses and ranavirus infection using a *Xenopus laevis* skin epithelial-like cell line

**DOI:** 10.1101/2021.06.29.450442

**Authors:** Lauren A. Todd, Maxwell P. Bui-Marinos, Barbara A. Katzenback

**Affiliations:** Department of Biology, Faculty of Science, University of Waterloo, 200 University Avenue West, Waterloo, Ontario, Canada, N2L 3G1

**Keywords:** Frog virus 3, microRNAs, antiviral immunity, poly(I:C), skin epithelial cells, *Xenopus laevis*

## Abstract

Epigenetic regulators such as microRNAs are emerging as conserved regulators of innate antiviral immunity in vertebrates, yet their roles in amphibian antiviral responses remain uncharacterized. We profiled changes in microRNA expressions in the *Xenopus laevis* skin epithelial-like cell line Xela DS2 in response to poly(I:C) – an analogue of double-stranded viral RNA and inducer of type I interferons – or frog virus 3 (FV3), an immunoevasive virus associated with amphibian mortality events. We sequenced small RNA libraries generated from untreated, poly(I:C)-treated, and FV3-infected cells. We detected 136 known *X. laevis* microRNAs and discovered 133 novel *X. laevis* microRNAs. Sixty-five microRNAs were differentially expressed in response to poly(I:C), many of which were predicted to target regulators of antiviral pathways such as cGAS-STING, RIG-I/MDA-5, TLR signaling, and type I interferon signaling, as well as products of these pathways (NF-κB-induced and interferon-stimulated genes). In contrast, only 49 microRNAs were altered by FV3 infection, fewer of which were predicted to interact with antiviral pathways. Interestingly, poly(I:C) treatment or FV3 infection downregulated transcripts encoding factors of the host microRNA biogenesis pathway. Our study is the first to suggest that host microRNAs regulate innate antiviral immunity in frogs, and sheds light on microRNA-mediated mechanisms of immunoevasion by FV3.

## Introduction

Epigenetic mechanisms are emerging as key regulators of immune signaling in vertebrates, and microRNAs (miRNAs) are a prime example of such a mechanism (Boosani and Agrawal 2016). miRNAs are a class of small non-coding RNAs that function in post-transcriptional regulation of gene expression. miRNA genes are transcribed in the nucleus by RNA polymerase II and are processed into hairpin precursors (pre-miRNAs) by the RNase III enzyme Drosha (Lee et al. 2002). Pre-miRNAs are exported to the cytoplasm where they are further processed into mature miRNA duplexes by the RNase III enzyme Dicer (Lee et al. 2002). One strand of the miRNA duplex is degraded, while the other guides the Argonaute-containing RNA-induced silencing complex (RISC) to mRNAs which are complementary to the miRNA guide (Lee et al. 2002). Direct interactions between the miRNA and mRNA [typically the 3’ untranslated region (UTR)] induce degradation or inhibit translation of the mRNA (Lee et al. 2002).

Vertebrate miRNAs have been found to regulate the expression of genes involved in antiviral signaling pathways that produce interferons (IFNs) and inflammatory cytokines in response to pathogen infection (Boosani and Agrawal 2016). Host miRNAs have been shown to regulate components of key pathogen sensing pathways such as the cytosolic RIG-I/MDA5 sensing pathway (Li et al. 2020), endosomal TLR3 pathway (Tili et al. 2007; Hou et al. 2009; Imaizumi et al. 2010), endosomal/cell surface TLR4 pathway (Wendlandt et al. 2012), and endosomal TLR7/8/9 pathways (Hou et al. 2009; Tang et al. 2009), as well as the JAK-STAT signaling pathway that regulates the production of IFN-stimulated genes (ISGs) (Tang et al. 2009; Jarret et al. 2016). In some cases, these interactions promote efficient host immune responses. However, viral infection can induce changes in host miRNA expression that detriment the host. For example, Coxsackievirus B3-induced miR-30a represses TRIM25 and RIG-I function, resulting in enhanced viral replication (Li et al. 2020), vesicular stomatitis virus-induced miR-146a represses type I IFN production and promotes viral replication by repressing TRAF6, IRAK2, and IRAK1 (Hou et al. 2009), and hepatitis C virus upregulates miR-208b and miR-499a-5p which enhances viral replication by dampening type I IFN signaling through repression of the type I IFN receptor IFNAR1 (Jarret et al. 2016). miRNA-mediated regulation of host immune responses is therefore complex, and while a growing number of functional studies have been conducted, our understanding of the broad functions host miRNAs play in antiviral responses remains limited.

Ranaviruses, type species frog virus 3 (FV3), are large double-stranded DNA viruses causing emerging infectious diseases that threaten amphibian populations. Despite the devastating amphibian morbidities and mortalities associated with ranaviral infection, our understanding of frog antiviral defenses remains in its infancy, and the roles of frog miRNAs in regulating these antiviral responses are largely unexplored. As frog skin is an important barrier to pathogen entry (Varga et al. 2019), we have recently generated a *Xenopus laevis* dorsal skin epithelial-like cell line (Xela DS2) (Bui-Marinos et al. 2020) that is permissive to FV3 (Bui-Marinos et al. submitted) to facilitate our understanding of antiviral responses in frog skin epithelial cells. All viruses, including FV3 (Doherty et al. 2016), produce viral double-stranded RNA (dsRNA) at some point in their replication. Along with others, we have studied frog cell antiviral responses through treatment with poly(I:C) (Sang et al. 2016; Wendel et al. 2018; Bui-Marinos et al. 2020), a synthetic viral dsRNA analog and known inducer of type I IFNs. While poly(I:C) is not a perfect mimic of viral dsRNA, as the sequence and length of viral dsRNA is known to impact the induction of host antiviral responses (Poynter and DeWitte-Orr 2015), it permits comparisons of type I IFN responses across cell types and species. By modeling “typical” antiviral responses in frogs using poly(I:C), we can further our understanding of how miRNAs function to regulate effective antiviral responses. By comparing “typical” antiviral miRNA responses in frog skin epithelial cells [modeled by poly(I:C)] to antiviral miRNA responses to immunoevasive viruses such as FV3, we aim to elucidate how viral pathogens may subvert epigenetic regulatory responses as direct or indirect mechanisms of immunoevasion.

In this study, we sought to perform initial investigations into the role of host miRNAs in innate antiviral immune responses in frog skin epithelial cells. The goals of this study were to (1) profile changes in miRNA expression in Xela DS2 skin epithelial-like cells during antiviral responses to poly(I:C) and FV3, (2) determine the antiviral targets of differentially expressed miRNAs, (3) compare normal antiviral miRNA responses modeled by poly(I:C) – a known inducer of robust antiviral responses – to miRNA responses induced by FV3, an immunoevasive virus, and (4) expand the number of currently annotated frog miRNAs through the discovery of novel *X. laevis* miRNAs.

## Materials and methods

### Cell line maintenance

Xela DS2, a skin epithelial-like cell line previously generated from *X. laevis* dorsal skin (Bui-Marinos et al. 2020), was maintained in seven parts Leibovitz’s L-15 (AL-15) medium (Wisent, Mont-Saint-Hilaire, Canada) diluted with 3 parts sterile cell culture water to adjust for amphibian osmolarity, and is herein referred to as amphibian L-15 (AL-15) medium. AL-15 medium was supplemented with 15% fetal bovine serum (FBS; lot #234K18; VWR, Radnor, United States) and used to culture Xela DS2 cells at 26 °C in 75 cm^2^ plug-seal tissue-culture treated flasks (BioLite, Thermo Fisher Scientific, Waltham, United States). Cells were split 1:4 upon reaching ~90-95% confluency, which usually occurred every 3 - 4 days. Epithelioma Papulosum Cyprini (EPC) cells were maintained in Leibovitz’s L-15 medium supplemented with 10% FBS. EPC cells were cultured in 75 cm^2^ plug-seal tissue-culture treated flasks at 26 °C and were split 1:4 weekly. Prior to use in experiments, viable Xela DS2 and EPC cells we enumerated using Trypan blue (0.2% final concentration; Invitrogen, Waltham, United States) and a haemocytometer, and seeded to account for their respective plating efficiencies (79% for Xela DS2 and 100% for EPC).

### Propagation of FV3 and determination of FV3 viral titres

FV3 (Granoff strain) was propagated in EPC cells as described previously (Bui-Marinos et al. submitted). Briefly, a monolayer of EPC cells was infected with a 1:10 dilution of stock FV3 in L-15 medium supplemented with 2% FBS for seven days at 26 °C. Seven days post-infection, virus-containing cell culture medium was collected, centrifuged at 1,000 × *g* for 10 min, and filtered through a 0.22 μm PES filter (FroggaBio, Concord, Canada) prior to storage at −80 °C until use.

The tissue culture infectious dose wherein 50% of cells are infected (TCID_50_/mL) values were determined according to the modified Kärber method (Kärber 1931; Pham et al. 2011). EPC cells (100,000/well) were seeded in a 96-well plate (BioBasic, Toronto, Canada) and allowed to adhere overnight at 26 °C. The following day, cell culture medium was removed, and cells were treated with 200 μL of a ten-fold dilution series of the virus-containing cell culture supernatants to be tested (FV3 stock prepared as described above or experimental sample supernatant) diluted in L-15 medium containing 2% FBS. Plates were sealed with parafilm and incubated at 26 °C for seven days prior to visual examination of cytopathic effects (CPE). EPC monoloayers were scored for CPE and approximate FV3 plaque forming unit (PFU)/mL values were calculated by multiplying TCID_50_/mL values by a factor of 0.7 (Knudson and Tinsley 1974). PFU/mL values were used to calculate MOI (Knudson and Tinsley 1974).

### Exposure of Xela DS2 to poly(I:C) or FV3

Xela DS2 cells (passages 77, 80, and 83; *n* = 3) were seeded into two 6-well plates (700,000 cells/well) in AL-15 medium supplemented with 15% FBS and allowed to form a monolayer overnight at 26 °C. The following day, cell culture medium was removed, and cells were treated with 2 mL of medium alone (AL-15 supplemented with 2% FBS, two wells per plate) or medium containing FV3 at an MOI of 2 (FV3 infected, one well per plate). After 2 h of incubation at 26 °C, the medium was removed, cells were washed three times with 2 mL amphibian phosphate buffered saline (APBS; 8 g/L sodium chloride, 0.5 g/L potassium chloride, 2.68 g/L sodium phosphate dibasic heptahydrate, 0.24 g/L potassium phosphate monobasic), and 2 mL of fresh AL-15 medium supplemented with 2% FBS was added to all wells. At this point, 1 μg/mL of poly(I:C) (Sigma-Aldrich, Burlington, United States) was added to one of the untreated wells. Untreated, poly(I:C)-treated, and FV3-infected cells were incubated at 26 °C for 24 h, while another set of untreated and FV3-infected cells were incubated at 26 °C for 72 h. Prior to collection of medium to determine TCID_50_/mL values and cells for RNA extraction, cell images were captured using a Leica DMi1 phase-contrast microscope fitted with an MC170 colour camera and LASX 4.8 software.

### RNA isolation

Adherent Xela DS2 cells and any detached cells in suspension were harvested for total RNA isolation using the Monarch Total RNA Minipreps kit (New England Biolabs, Ipswich, United States) according to the manufacturer’s recommended protocol. Suspension cells present in the cell culture medium were collected by centrifugation at 500 *× g* for 2 min and the resulting cell culture medium was aspirated. Lysis buffer (800 μL) was added to each well and incubated with adherent cells for 2 min at room temperature. Lysed cells were transferred from the 6-well plates to microcentrifuge tubes containing the corresponding pelleted suspension cells. Cells were resuspended by pipetting and incubated for 2 min to permit cell lysis. At this stage, an exogenous miRNA (cel-miR-39 Spike-In kit; Norgen Biotek Corp., Thorold, Canada) was spiked in (99 fmol) to serve as a positive control for miRNA detection. On-column DNase I treatment was performed for enzymatic removal of residual genomic DNA according to the manufacturer’s instructions, and elution was performed in nuclease-free water (50 μL). Total RNA was quantified using the Take3 microvolume plate accessory on a Cytation 5 Cell Imaging Multi-Mode Reader (Biotek, Winooski, United States), RNA purity was determined by analyzing A_260/280_ and A_260/230_ ratios, and RNA integrity was examined by electrophoresis on a bleach agarose gel (Aranda et al. 2012).

### Reverse transcription polymerase chain reaction (RT-PCR) detection of a control spike-in miRNA prior to small RNA-seq

For RT-PCR analysis of cel-miR-39 expression, 500 ng of DNase I-treated total RNA was reverse-transcribed into cDNA using the microScript miRNA cDNA Synthesis Kit (Norgen Biotek Corp., Thorold, Canada). Resulting cDNA was diluted four-fold and subjected to end-point PCR using Taq DNA Polymerase (GeneDirex Inc., Taoyuan, Taiwan) in a Mastercycler Nexus (Eppendorf, Hamburg, Germany) thermocycler. PCR reaction conditions were as follows: 1× GeneDirex buffer, 200 μM dNTPs, 0.2 μM forward and reverse primers, 0.625 U GeneDirex Taq, and 2 μL diluted cDNA. Spiked-in cel-miR-39 transcripts were amplified to confirm the ability to detect miRNAs, while *actb* transcripts were amplified as an internal control for amplifiability. PCR cycling conditions were as follows: 94 °C for 3 min, 40 × [94 °C for 15 s, 60 °C for 30 s, 72 °C for 45 s], and a final hold at 4 °C. All primers used in this study are listed in **Table S1**.

### Small RNA library preparation and sequencing

For each sample, ~2 μg of RNA (~80 ng/μL in 25 μL) was shipped to The Centre for Applied Genomics (TCAG) at The Hospital for Sick Children (Toronto, Canada) on dry ice. TCAG performed further quality assessment using a Bioanalyzer 2100 and RNA 6000 Nano LabChip Kit (Agilent Technologies Inc., Santa Clara, United States), and samples with an RNA integrity number (RIN) > 8.5 were used for small RNA-seq library preparation. Library preparation and sequencing was performed by TCAG. Small RNA-seq libraries were prepared using the NEBNext Multiplex Small RNA Library Prep Set for Illumina (New England Biolabs, Ipswich, United States). Small RNAs were enriched using bead-based size selection. Resulting small RNA libraries were sequenced on an Illumina HiSeq 2500 rapid-run flow cell to generate 50 nucleotide (nt) single-end reads.

### Small RNA-seq read pre-processing and mapping

Raw ~50 nt single-end sequencing reads were adaptor-trimmed and size-filtered using Cutadapt (Martin 2011) with the following parameters: single-end reads, 3’ adaptor of AGATCGGAAGAGCACACGTCTGAACTCCAGTCAC, minimum length of 17, maximum length of 30. Read quality reports were generated using FastQC (Andrews 2010) with default parameters. High quality (Q > 34; 0.04% error rate) clean reads were mapped to the *X. laevis* genome (J strain; GCF_001663 975.1) using Bowtie2 (Langmead and Salzberg 2012) with the following parameters: single-end reads, write aligned and unaligned reads to separate files, allow one mismatch, output in SAM format. Reads that mapped to the *X. laevis* genome were retained for further analyses. Cutadapt, FastQC, and Bowtie2 were run using the Galaxy web platform (usegalaxy.org; (Afgan et al. 2016)).

### Detection of novel X. laevis miRNAs

Novel *X. laevis* miRNAs were detected using miRDeep2 v0.1.3 (Friedländer et al. 2012). Aligned reads were collapsed into unique sequences using the mapper.pl script and reads with counts < 10 in a given library were discarded for this purpose. Transcripts with read counts < 10 are generally considered to be noise and are often removed prior to downstream bioinformatics analysis (Law et al. 2016), thus removing these reads prior to novel miRNA prediction helps improve the signal-to-noise ratio, which is an important component of miRDeep2-based miRNA detection. The miRDeep2.pl script was executed using default parameters and the following inputs: collapsed reads, alignments, the *X. laevis* genome, known *X. laevis* pre-miRNAs (miRBase), and known *X. laevis* and *Xenopus tropicalis* miRNAs (miRBase). Secondary structures associated with novel *X. laevis* pre-miRNAs were determined using the RNAFold component of the ViennaRNA package v2.4.17 (Lorenz et al. 2011) under default parameters. Novel *X. laevis* miRNAs were considered high confidence if they were characterized by a miRDeep2 score ≥ 4 (≥ 90% probability of being a true positive) or if their sequence perfectly matched a known *X. tropicalis* miRNA.

### Quantification and differential expression analysis of known and novel X. laevis miRNAs

*X. laevis* miRNA read counts were generated using miRDeep2 (Friedländer et al. 2012). Collapsed *X. laevis* reads, known and novel *X. laevis* miRNA sequences, and *X. laevis* pre-miRNA sequences [known (retrieved from miRBase) and novel (predicted by miRDeep2)] were input into the miRDeep2 quantifier.pl script using default parameters. Reads corresponding to several miRNAs were in very low relative abundance (e.g. < 10 in a given library). To improve the accuracy of our analyses by strengthening the signal-to-noise ratio (Law et al. 2016), miRNAs were considered detected if their read counts were at least 10 in at least one library. miRNAs with < 10 read counts in all libraries were considered undetected and were excluded from downstream analyses. Read counts served as input for differential expression analysis using DESeq2 (Love et al. 2014) and EdgeR (Robinson et al. 2010). DESeq2 parameters included “select datasets per level”, input data = count data, and visualizing results = yes. EdgeR parameters included “single count matrix”, use factor information file = yes, use gene annotation = no, normalization method = TMM. Known and novel *X. laevis* miRNAs were considered differentially expressed if an FDR (false discovery rate) < 0.05 was reached by both programs (consensus DE). Heatmaps were generated using heatmap2 with the following parameters: enable data clustering = no, coloring groups = blue to white to red. DESeq2, EdgeR, and heatmap2 were run using the Galaxy web platform (usegalaxy.org; (Afgan et al. 2016)).

### miRNA target prediction

*X. laevis* 3’ UTRs were retrieved from the UCSC Table Browser using the Xenbase trackhub (genome.ucsc.edu; genome assembly v9.2). FV3 coding sequences (cds) were retrieved from NCBI (Accession NC_005946.1). Prior to running target prediction analysis, the dinucleotide frequencies of *X. laevis* 3’UTRs and FV3 cds were calculated using the compseq function of EMBOSS v6.6.0 (Rice et al. 2000). Using these frequencies in target prediction analyses improves *p*-value accuracy (Rehmsmeier et al. 2004). The targets of known and novel *X. laevis* miRNAs were predicted using RNAcalibrate and RNAhybrid v2.1.2 (Rehmsmeier et al. 2004). RNAcalibrate was first used to estimate distribution parameters using default parameters (in addition to strict seed pairing at bases 2-7 of the miRNA and the dinucleotide frequencies generated by EMBOSS). Next, RNAhybrid was used to predict the *X. laevis* and FV3 mRNA targets of known and novel *X. laevis* miRNAs. RNAhybrid parameters included maximum energy = −25 kcal/mol, maximum mismatches in internal loop = 1, strict seed pairing at miRNA bases 2-7, mismatches in bulge loop = 0, one hit per interaction, *p* < 0.01. Gene functions were annotated using Xenbase (Karimi et al. 2018). Immune-related *X. laevis* targets were identified through a combination of manual search and cross-referencing The Gene Ontology (GO) Resource (The Gene Ontology Consortium 2019) and InnateDB (Breuer et al. 2013).

### GO analysis

The PANTHER overrepresentation test was used to perform GO term enrichment analysis to reveal biological processes that are overrepresented in the list of *X. laevis* genes predicted to be targeted by differentially expressed *X. laevis* miRNAs (The Gene Ontology Consortium 2019). PANTHER parameters were as follows: reference list: all genes in database, “GO biological process complete” annotation dataset, Fisher’s exact test, FDR < 0.05. GO term analysis was performed at geneontology.org.

### Reverse transcription quantitative polymerase chain reaction (RT-qPCR)

RT-qPCR experiments were performed on a duplicate set of RNA samples isolated from cells treated in parallel with RNA-seq samples. Total RNA (1 μg) was reverse-transcribed using the microScript miRNA cDNA synthesis kit (Norgen Biotech Corp., Thorold, Canada) according to the manufacturer’s recommended protocol. Resulting cDNA was diluted (1/20) with nuclease-free water and stored at −20 °C until RT-qPCR analysis. RT-qPCR reactions consisted of 1 × PowerUP SYBR green mix (Thermo Fisher Scientific, Waltham, United States), 2.5 μL of diluted cDNA, and 500 nM (each) forward (miRNA-specific) and reverse (universal) primers in a final reaction volume of 10 μL. RT-qPCR primer sequences and efficiencies are presented in **Table S1**. All RT-qPCR reactions were performed in triplicate on a QuantStudio5 Real-Time PCR System (Thermo Fisher Scientific, Waltham, United States), and reaction conditions were as follows: denaturation at 50 °C for 2 min and 95°C for 2 min, followed by 40 cycles of denaturation at 95 °C for 1 s and extension at 60 °C for 30 s. A melt curve step (denaturation at 95 °C for 1 s and dissociation analysis at 60 °C for 20 s followed by 0.1 °C increments from 60 °C to 95 °C at 0.1 °C/s) was used to ensure the presence of a single dissociation peak.

RT-qPCR data was analyzed using the ΔΔCt method in Microsoft Excel with xla-miR-16a-5p serving as an endogenous control. xla-miR-16a-5p was chosen as a control for RT-qPCR normalization by examination of the small RNA-seq data [normalized reads per million (RPM) read counts generated by the miRDeep2 quantifier.pl script]. This control miRNA was chosen based on the following criteria: (1) < 10% variability in expression (relative to total number of host-mapped reads) between experimental libraries and their respective control libraries, and (2) > 1,000 RPM in the libraries of at least one treatment. xla-miR-16a-5p served as the endogenous control for RT-qPCRs targeting miRNAs, as well as *dicer1* and *drosha* transcripts. While we recognize that normalization of an mRNA to a miRNA control is not ideal, it is difficult to identify an appropriate endogenous control gene for use in infection experiments, particularly late in viral infection. As the relative proportion of host RNA in total RNA samples decreases as viral RNA levels increase in infected cells, it is difficult to experimentally validate an endogenous control gene by RT-qPCR alone, due to the inherent instability of host RNA abundance between untreated and infected cells. Therefore, we relied on our RNA-seq data to determine an appropriate endogenous control miRNA in lieu of an mRNA control gene, as we were able to accurately assess the stability of the control miRNA relative to the total host RNA abundance in each sample. Nevertheless, statistically significant differences were identified using one-way ANOVA tests paired with Dunnet’s post-hoc tests or Student’s *t*-tests where appropriate, and differences were considered statistically significant if *p* < 0.05.

### Data availability

The small RNA-seq datasets generated by this study will be available in the NCBI Short Read Archive (SRA) repository (accession number pending).

## Results

### Detection of X. laevis miRNAs in small RNA-seq libraries

To identify miRNAs that are involved in innate immune responses in frog skin epithelial cells, we sequenced small RNAs from untreated (24 h and 72 h), poly(I:C)-treated (24 h), and FV3-infected (24 h and 72 h) Xela DS2 skin epithelial-like cells. We chose a poly(I:C) concentration of 1 μg/mL and a treatment duration of 24 h as these conditions produce robust transcriptional immune responses in Xela DS2 cells that confer functional protection against FV3 infection (Bui-Marinos et al. submitted) and have been used in previous studies profiling miRNA responses to poly(I:C) treatment in other species (Wang et al. 2017; Wu et al. 2019). We chose FV3 sampling time points of 24 h and 72 h as overt FV3-induced CPE are not obvious at earlier time points (e.g. 12 h), moderate CPE is observed at 24 h, and extensive CPE is evident at 72 h in Xela DS2 cells infected with FV3 at a multiplicity of infection (MOI) of 2 (**Fig. S1a**). We experimentally determined that 72 h approaches, but falls short of, the threshold where FV3-infected Xela DS2 cells are too compromised to obtain RNA of sufficient quality and quantity (96 h; data not shown). Small RNAs were sequenced on an Illumina HiSeq 2500 resulting in 15 small RNA libraries (*n* = 3 per treatment; **Fig. S1b-d**). A schematic of the bioinformatics pipeline employed in this study is depicted in **Fig. 1a**. Approximately 7.6-11.3 million raw reads per sample underwent adapter, length, and quality filtering using Cutadapt (Martin 2011) and FastQC (Andrews 2010) to obtain ~1.5-5.9 million clean reads (17-30 nt) per library (**Figure 1b**). Genome alignment was performed using Bowtie2 (Langmead and Salzberg 2012). Reads that aligned to the *X. laevis* genome were subjected to further analyses. The number of *X. laevis*-mapped reads in each library ranged from ~29% to 97% (**Fig. 1b**) and their size distribution spanned from 17 nt to 30 nt (**Fig. 1c**). More than 79% of the small RNA reads that mapped to the *X. laevis* genome were 21-24 nt in length, with 22-23 nt RNAs representing the majority.

**Fig. 1.**
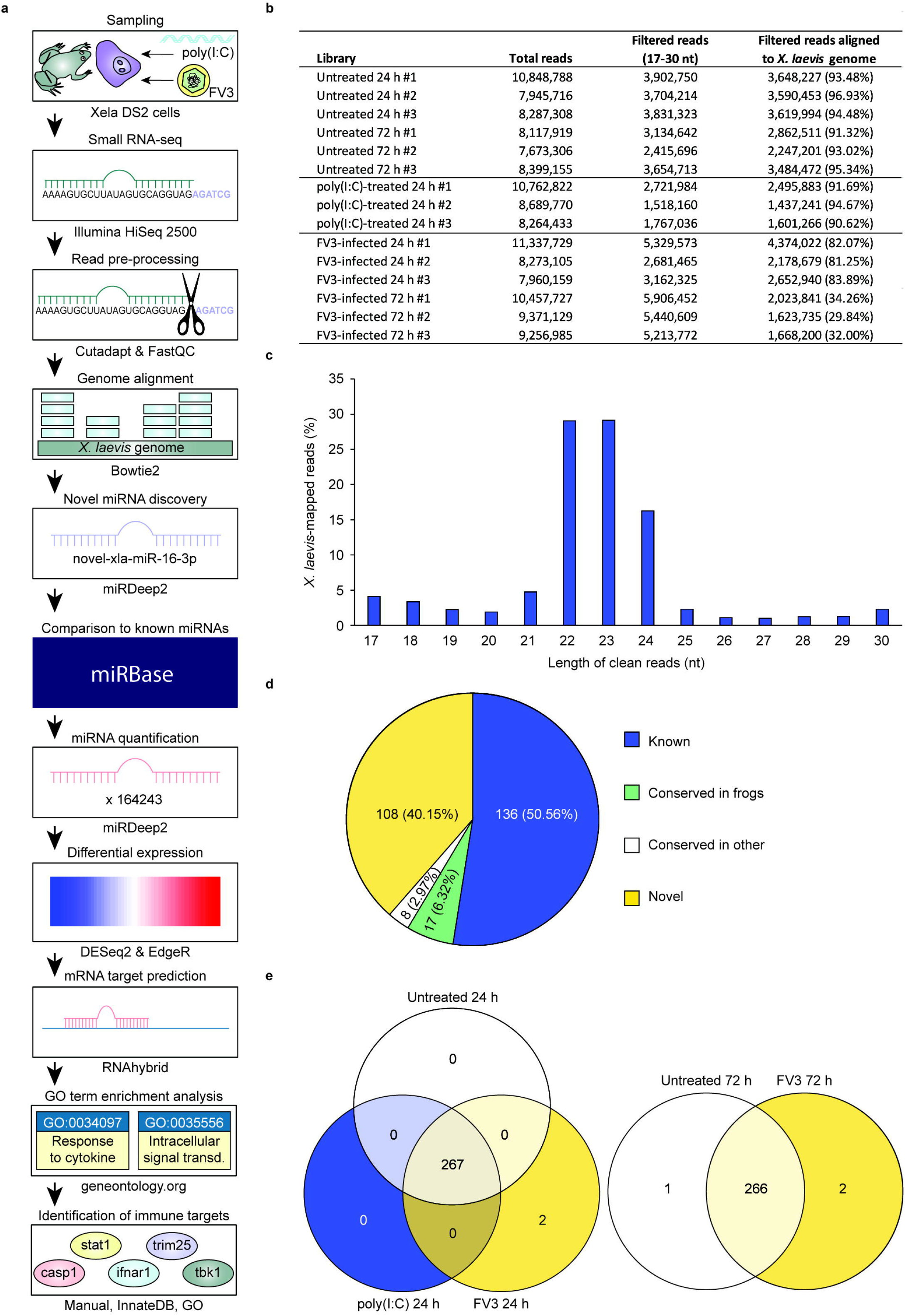
Characteristics of *X. laevis* small RNA-seq libraries. (**a**) A schematic representation of the bioinformatics pipeline used in this study. (**b**) The total number of small RNA reads, the number of reads passing size filtering (17-30 nt), and the number of filtered reads that aligned to the *X. laevis* genome for all 15 small RNA libraries. Library numbers correspond to the respective independent experiment. (**c**) The size distribution of *X. laevis-mapped* reads across all libraries. (**d**) Different categories of miRNAs detected in Xela DS2 cells by small RNA-seq. “Known” miRNAs refer to *X. laevis* miRNAs present in miRBase, “conserved in frogs” refer to miRNA orthologs of known *X. tropicalis* miRNAs in miRBase not yet reported in *X. laevis*, “conserved in other” miRNAs refer to orthologs of miRNAs in miRBase not yet reported in frogs, and “novel” miRNAs refer to new miRNAs that are not present in miRBase in any species. (**e**) Venn diagram of the *X. laevis* miRNAs detected in small RNA-seq libraries generated from untreated, poly(I:C)-treated, and FV3-infected Xela DS2 cells (left = 24 h and right = 72 h).

At present, there are 247 mature *X. laevis* miRNAs in the miRBase database (Kozomara et al. 2019), a number which is relatively low for vertebrates. Thus, we sought to first identify novel *X. laevis* miRNAs from our small RNA libraries. Using miRDeep2 software, we detected 133 novel *X. laevis* miRNAs which we classified into 52 families (**Table S2**). Twenty-five of these novel *X. laevis* miRNAs are conserved in other species and present in miRBase, while the remainder are entirely novel miRNAs in any species (**Fig. 1d**). Many of these entirely novel miRNAs are derived from repetitive genomic regions (**Table S2**). All 133 novel *X. laevis* miRNAs will be submitted to miRBase upon manuscript acceptance (Kozomara et al. 2019). Conserved miRNAs were assigned the name of the corresponding orthologous miRNA in miRBase, while truly novel miRNAs were numbered sequentially beginning with the next available number in miRBase (Kozomara et al. 2019). There is a bias towards “U” as the first (5’) nucleotide of these novel *X. laevis* miRNAs, which is characteristic of vertebrate miRNAs (Yang et al. 2011). Next, we used miRDeep2 software to detect and quantify known and novel *X. laevis* miRNAs in each library. Along with the 133 novel *X. laevis* miRNAs, we detected 136 known *X. laevis* miRNAs across all 15 libraries (**Fig. 1d**). Thus, we detected a total of 269 *X. laevis* miRNAs in our small RNA-seq datasets. The majority of these miRNAs were shared amongst all groups, however two miRNAs were uniquely detected in FV3-infected cells at 24 h and 72 h, while one miRNA was uniquely detected in untreated cells at 72 h (**Fig. 1e**).

### X. laevis miRNAs are differentially expressed in response to poly(I:C) and target important regulators of antiviral immunity

To begin elucidating the importance of frog miRNAs to innate antiviral responses, we profiled changes in *X. laevis* miRNA expression in response to the viral dsRNA analogue poly(I:C). We chose poly(I:C) to model a “typical” antiviral response in frog skin epithelial cells as poly(I:C) is a potent inducer of type I IFNs and downstream antiviral responses in Xela DS2 cells (Bui-Marinos et al. 2020) and in other vertebrate systems from humans to fish (Tamassia et al. 2008; Poynter and DeWitte-Orr 2015). Furthermore, prior treatment of Xela DS2 with low doses of poly(I:C) induces functional protective antiviral responses against FV3 (Bui-Marinos et al. submitted). Thus, poly(I:C) represents an ideal molecule to use in modeling potent and functional antiviral responses involving miRNAs for comparison with FV3-induced responses, which may be less representative of an effective miRNA-mediated antiviral response given that FV3 is immunoevasive (Grayfer et al. 2015; Robert et al. 2017).

We performed differential expression analysis on the 269 (136 known and 133 novel) detected *X. laevis* miRNAs, comparing the 24 h untreated Xela DS2 libraries with poly(I:C)-treated Xela DS2 libraries. miRNAs were considered differentially expressed if differential expression was identified by both EdgeR and DESeq2 programs (consensus) and when FDR < 0.05. We discovered that 47 known *X. laevis* miRNAs are differentially expressed in response to poly(I:C) (23 downregulated and 24 upregulated) (**Fig. 2a**), while 18 novel *X. laevis* miRNAs are differentially expressed in response to poly(I:C) (seven downregulated and 11 upregulated) (**Fig. 2b**). These 65 miRNAs may represent key post-transcriptional mediators of effective innate antiviral responses in frog skin epithelial cells.

**Fig. 2.**
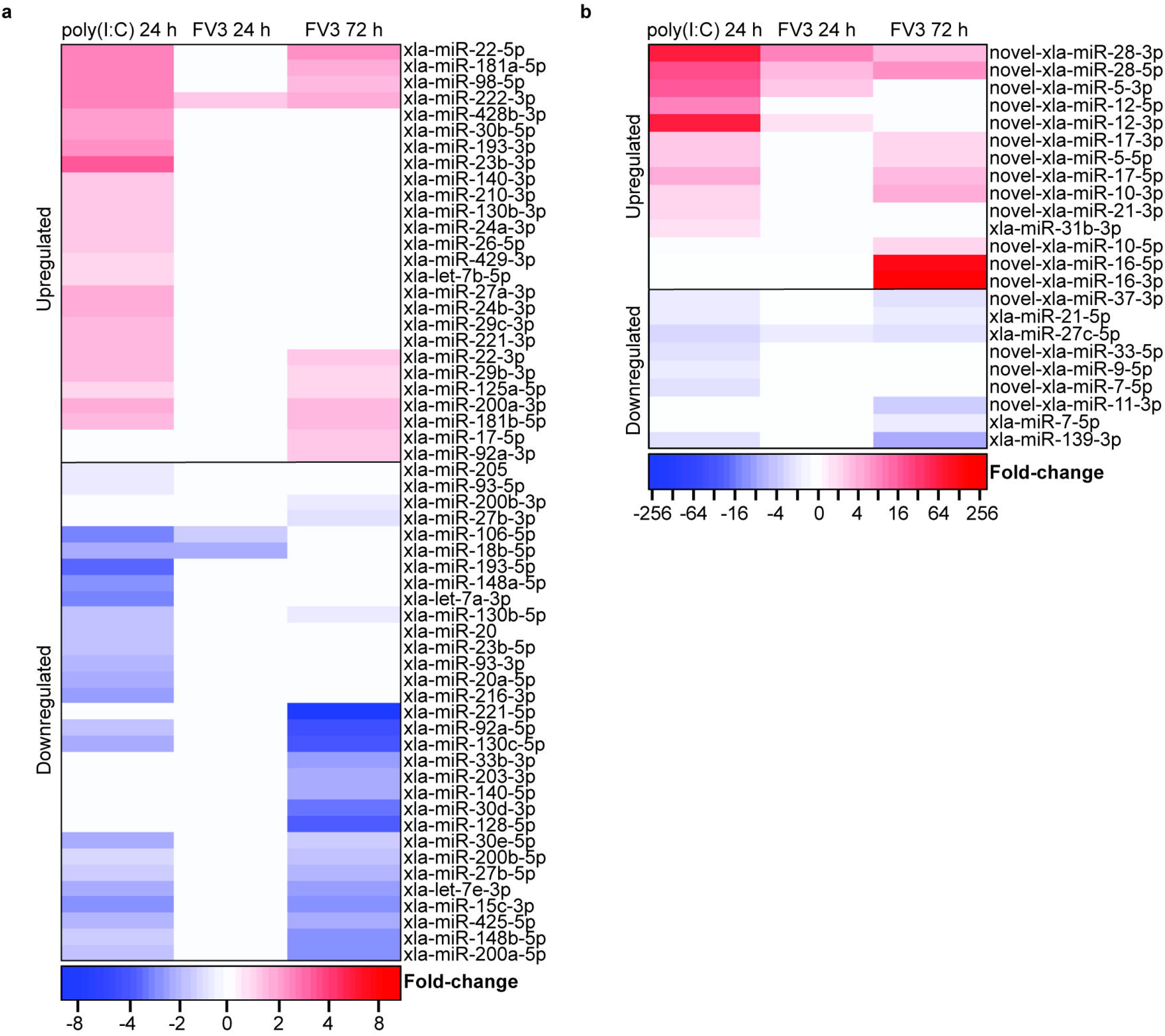
Differential expression analysis of *X. laevis* miRNAs in response to poly(I:C) or FV3. Identification of known (**a**) and novel (**b**) *X. laevis* miRNAs that are differentially expressed in Xela DS2 cells in response to poly(I:C) or FV3 (compared to untreated cells). Statistically significant (FDR < 0.05, *n* = 3, Wald test) decreases and increases in miRNA levels relative to corresponding untreated controls are represented by blue and red, respectively. The indicated fold-change values were determined by DESeq2 but were also statistically supported by EdgeR. White bars represent relative fold-changes that were not statistically significant.

In order to begin identifying the potential functions of *X. laevis* miRNAs that are differentially expressed in response to poly(I:C), we performed target prediction analyses using RNAhybrid (Rehmsmeier et al. 2004) to predict their host gene targets. We predicted robust interactions between the 65 *X. laevis* miRNAs that are differentially expressed in response to poly(I:C) and the 3’ UTRs of 2,066 endogenous host genes (1,937 if consolidating the L and S homeologs) (**Table S4**). GO analysis of the predicted *X. laevis* targets of miRNAs that are differentially expressed in response to poly(I:C) revealed statistical enrichment of numerous functional categories, including many related to antiviral defenses such as “negative regulation of I-kappaB kinase/NF-kappaB signaling”, “transforming growth factor beta receptor signaling pathway”, “viral process”, “intracellular signal transduction”, “cellular response to stress”, and “regulation of apoptotic process” (**Fig. 3a**).

**Fig. 3.**
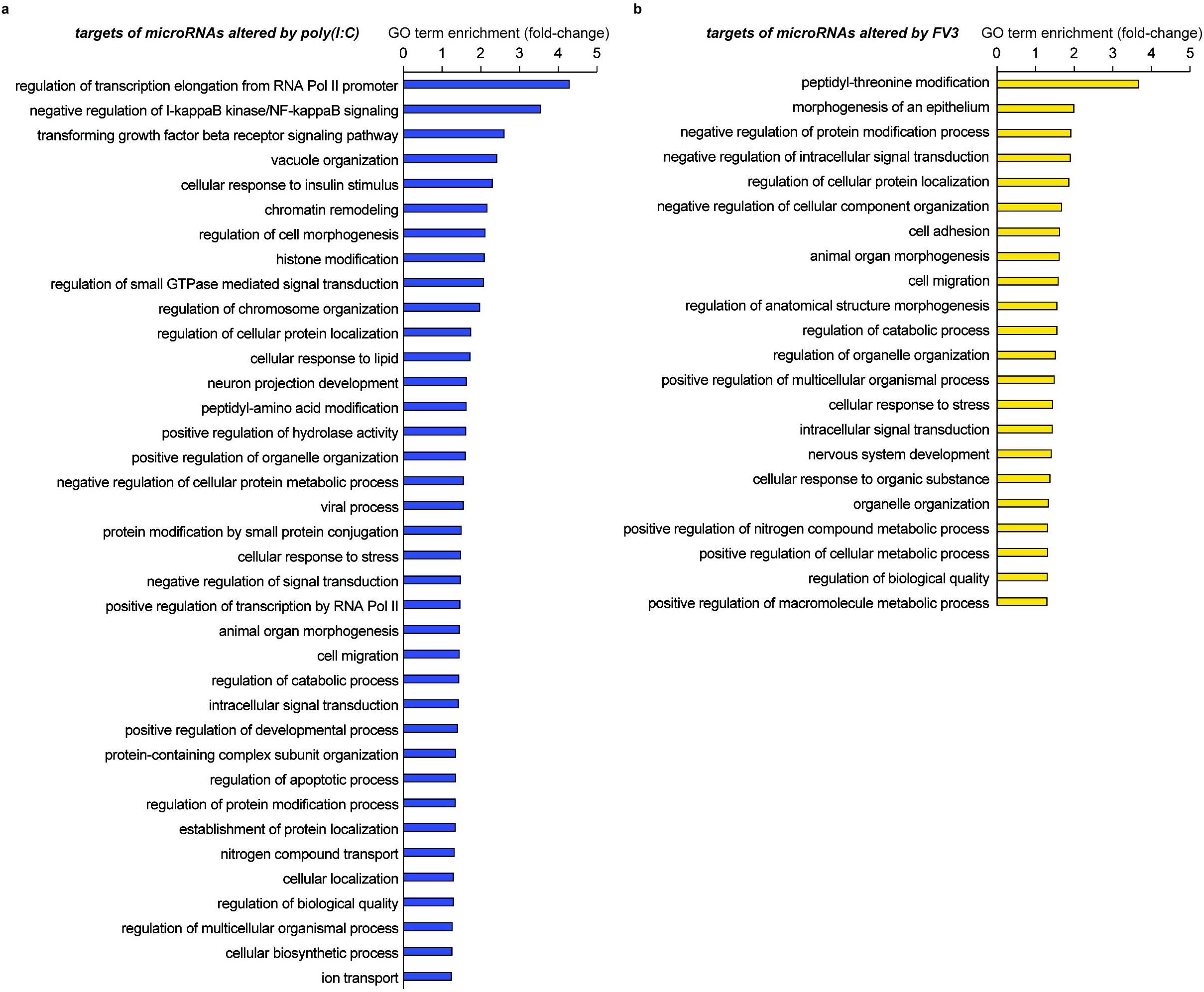
GO term enrichment analysis of *X. laevis* 3’ UTRs predicted to be targets of differentially expressed *X. laevis* miRNAs. GO terms enriched in the list of targets of miRNAs with altered expression in response to poly(I:C) (**a**) or FV3 (**b**). In cases where multiple GO terms from the same hierarchy were found to be enriched, the most specific GO term is listed. An FDR value of less than 0.05 was considered statistically significant (Fisher’s exact test).

### X. laevis miRNAs are differentially expressed in response to FV3 and target important regulators of antiviral immunity

We have recently demonstrated that Xela DS2 cells are susceptible and permissive to FV3 but fail to induce antiviral programs in response to FV3 (Bui-Marinos et al. submitted). As FV3 is believed to be immunoevasive and is known to produce dsRNA during its replication (Doherty et al. 2016), we sought to compare host miRNA responses to FV3 with host miRNA responses to poly(I:C) to develop an understanding of how FV3-induced miRNA responses differ from typical antiviral responses mounted in response to viral dsRNA. We profiled changes in *X. laevis* miRNA expression in response to FV3 infection and uncovered a total of 49 miRNAs that were differentially expressed in response to FV3 at either time-point (24 h or 72 h). Three known miRNAs are differentially expressed in response to FV3 infection at 24 h (two downregulated and one upregulated), and 30 are differentially expressed in response to FV3 infection at 72 h (19 downregulated and 11 upregulated) (**Fig. 2a**). Five novel miRNAs are differentially expressed in response to FV3 infection at 24 h (one downregulated and four upregulated), and 15 are differentially expressed in response to FV3 infection at 72 h (six downregulated and nine upregulated) (**Fig. 2b**). While 34 of these differentially expressed miRNAs are also altered by poly(I:C) treatment, 15 are uniquely altered by FV3 infection.

To elucidate the targets of miRNAs that are differentially expressed in response to FV3, we performed target predictions as described in the previous section, but also considered potential viral targets of these miRNAs. We predicted robust interactions between five FV3 gene transcripts and five *X. laevis* miRNAs (three known and two novel) that are differentially expressed in response to FV3 (**Table S3**), suggesting that *X. laevis* miRNAs that are differentially expressed in response to FV3 infection may directly target FV3 viral transcripts. Additionally, we predicted robust interactions between the 3’ UTRs of 1,482 *X. laevis* genes (1,406 if consolidating the L and S homeologs) and 49 miRNAs that are differentially expressed in response to FV3 (**Table S4**). These findings suggest that 49 *X. laevis* miRNAs may play a role in regulating endogenous gene expression in response to FV3. GO term enrichment analysis on the list of targets of miRNAs that are altered in response to FV3 revealed enrichment of several GO terms, a few of which are immune-related, such as “cellular response to stress”, “intracellular signal transduction”, and “cellular response to organic substance” (**Fig. 3b**). Overall, the miRNAs altered by FV3 infection were associated with fewer immune-related functional categories than miRNAs altered by poly(I:C).

### Mapping of differentially expressed miRNAs to antiviral pathways

Pathogens are sensed by a variety of pattern recognition receptors (PRRs) that are either cytosolic (e.g. cGAS, RIG-I/MDA5) or membrane-associated [e.g. toll-like receptors (TLRs)] and recognize pathogen-associated molecular patterns (PAMPs) such as viral nucleic acids (Mogensen 2009; Motwani et al. 2019). PAMP sensing by PRRs can initiate antiviral signaling cascades that culminate in the production of IFNs and inflammatory cytokines that combat infection (Mogensen 2009). Thus, to gain a clearer picture of the antiviral pathways targeted by miRNAs that are differentially expressed in response to poly(I:C) or FV3, we mapped these miRNAs onto several key antiviral pathways such as cGAS-STING, RIG-I/MDA5, TLR3, TLR4, TLR7, TLR8, TLR9, and type I IFN signaling pathways (**Fig. 4**). Poly(I:C)-altered miRNAs target components of all eight of these pathways (*stim1, trim25, tbk1, tram1, ripk1, tab2, irak2, irf5, nmi, ifnar1, stat1, ubr4*, and *pkc-δ*), as well as their downstream products such as NF-κB-induced genes and ISGs (**Fig. 4a**). On the other hand, FV3-altered miRNAs targeted fewer components of these pathways (*trim14, stim1, trim25, tbk1, tram1, ripk1, tab2, stat1, ubr4*, and *pkc-δ*) and fewer NF-κB-induced genes and ISGs (**Fig. 4b**). While some targets in these pathways are found in common between miRNAs altered by poly(I:C) and FV3, there are several distinct differences. Poly(I:C)-altered miRNAs uniquely target *irak2, irf5, nmi*, and *ifnar1*, while FV3-altered miRNAs uniquely target *trim14*. Interestingly, while several poly(I:C)-altered miRNAs target components of TLR7/8/9 signaling, FV3 is associated with a lack of changes in the expression of miRNAs targeting these pathways.

**Fig. 4.**
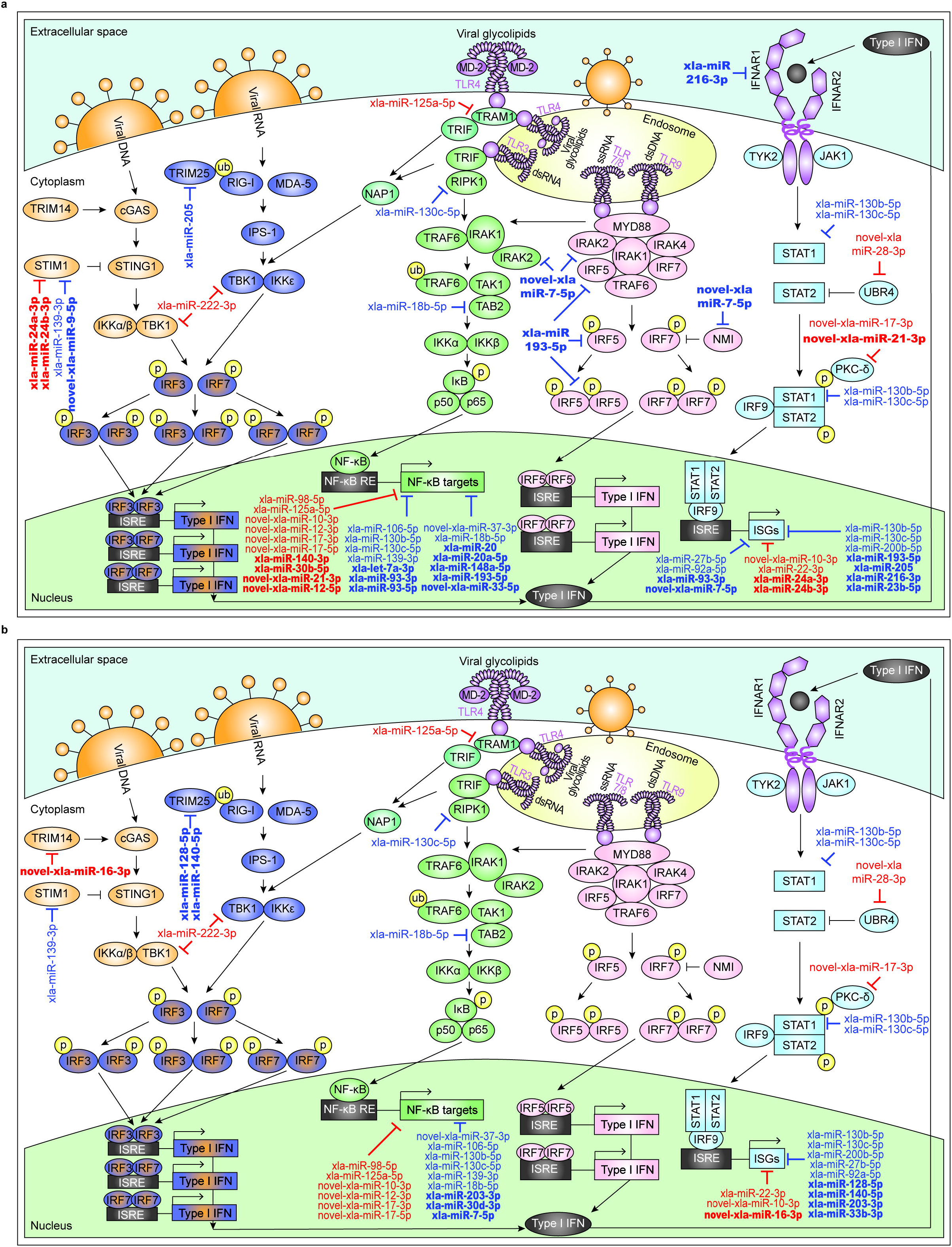
Differentially expressed *X. laevis* miRNAs are predicted to target important regulators of innate antiviral immunity. Depicted are miRNAs that are predicted to target key genes in innate antiviral pathways and are differentially expressed in response to poly(I:C) (**a**) or FV3 (**b**). Upregulated miRNAs are in red and downregulated miRNAs are in blue. miRNAs with differential expression unique to either treatment are bolded. Pathway schematics were adapted from (InvivoGen). Note: IRF3 is produced and phosphorylated, and its activity stimulates type I IFN production (JAK-STAT pathway), which stimulates the production of IRF7 as an ISG (Marie et al. 1998; Sato et al. 1998). IRF7 is then phosphorylated and homodimerizes or heterodimerizes with phosphorylated IRF3 to enhance type I IFN production (Marie et al. 1998; Sato et al. 1998; Honda et al. 2006). dsDNA = double-stranded DNA; dsRNA = double-stranded RNA; IFN = interferon; ISGs = IFN-stimulated genes; ISRE = IFN-stimulated response element; NF-κB RE = NF-κB response element; p = phosphorylation; ssRNA = single-stranded RNA; ub = ubiquitination.

### Validation of key differentially expressed miRNAs that target immune-related genes

We validated the differential expression profiles of a subset of *X. laevis* miRNAs that are predicted to target important regulators of innate antiviral responses. We chose 12 miRNAs that (1) are predicted to target important immune-related genes, (2) represent a combination of known, conserved, and novel miRNAs, (3) represent a combination of upregulated and downregulated miRNAs, and (4) represent a combination of miRNAs with changes in expression that are consistent between the poly(I:C) and FV3 treatment groups or are inconsistent between these groups. These miRNAs included xla-miR-106-5p, xla-miR-130c-5p, xla-miR-139-3p, xla-miR-140-5p, xla-miR-222-3p, novel-xla-miR-7-5p, xla-miR-22-5p, novel-xla-miR-12-3p, novel-xla-miR-16-5p, novel-xla-miR-16-3p, novel-xla-miR-28-5p, and novel-xla-miR-28-3p. We successfully validated the RNA-seq-derived differential expression profiles of 11 of these miRNAs by RT-qPCR (**Fig. 5**). Although the magnitude of differential expression differed between the two techniques (RT-qPCR and RNA-seq), the overall trends remained the same. However, one miRNA (xla-miR-222-3p) did not show comparable differential expression profiles between the two techniques.

**Fig. 5.**
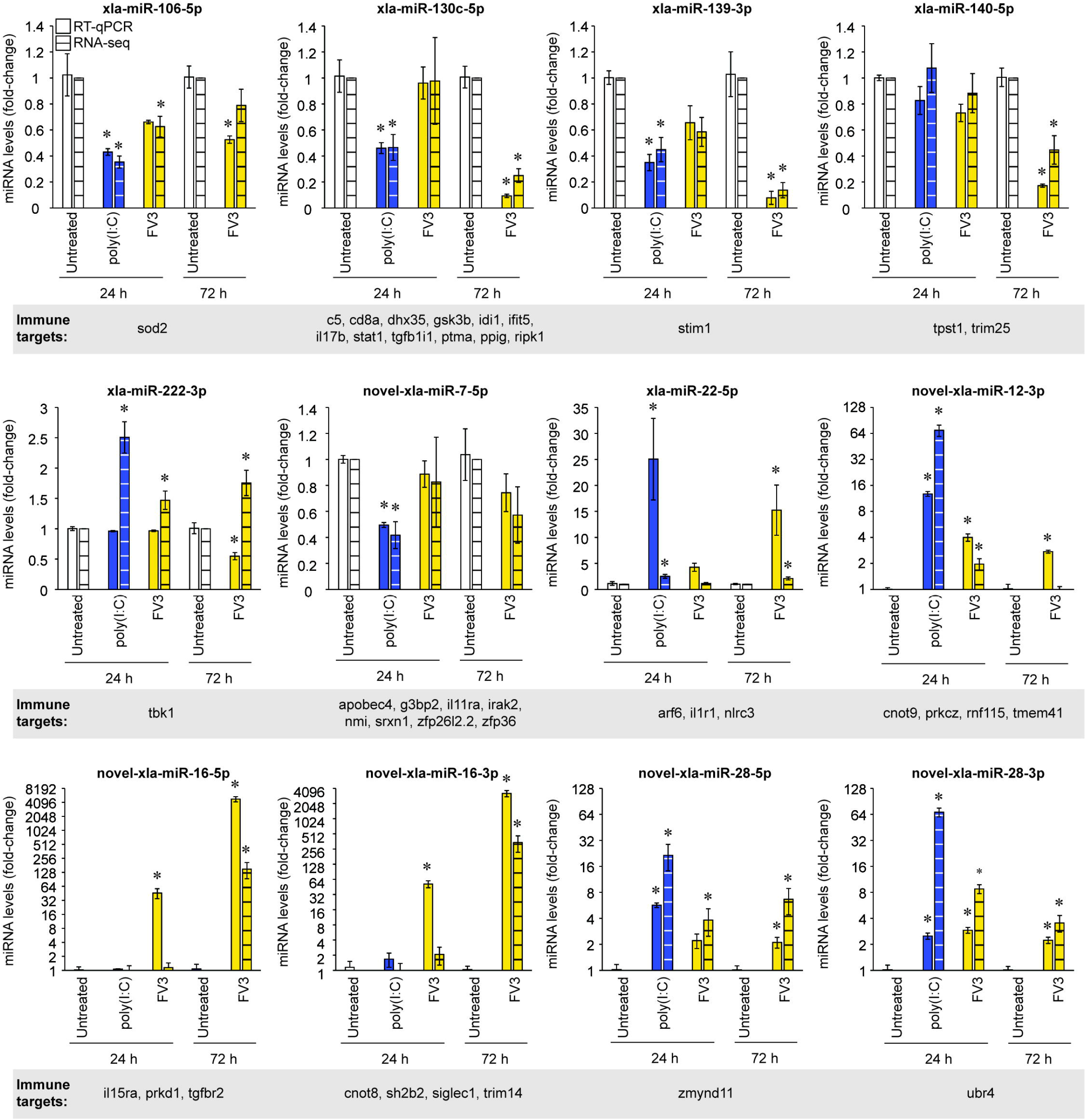
RT-qPCR validation of a subset differentially expressed *X. laevis* miRNAs. cDNA synthesized from total RNA isolated from untreated, poly(I:C)-treated and FV3-infected Xela DS2 cells was subjected to RT-qPCR analysis using forward primers specific to each miRNA of interest and a universal reverse primer. RT-qPCR results are depicted in solid colours and differential expression analysis of the RNA-seq data is depicted in striped corresponding colours for comparison. RT-qPCR data were analyzed using the ΔΔCt method using xla-miR-16a-5p (levels unchanged by any treatment based on RNA-seq data) as an internal control. Data is depicted as mean ± standard error (*n* = 3), and values corresponding to treatment groups are presented relative to their respective untreated control, which was set to a reference value of 1. Statistical significance was assessed using one-way analysis of variance (ANOVA) tests paired with Dunnett’s post-hoc tests (24 h) or unpaired Student’s *t*-tests (72 h), with a *p* value < 0.05 considered statistically significant from the time-matched untreated control (within the same miRNA detection method), as denoted by asterisks (*). Immune gene targets of each miRNA predicted using RNAhybrid are depicted below the corresponding graph.

### Poly(I:C) treatment downregulates dicer1 transcripts, while FV3 infection represses dicer1 and drosha transcripts

miRNAs that target immune-related genes were more frequently downregulated than upregulated in response to either poly(I:C) or FV3. In line with this, and as observed in other pathosystems, host cells have been found to modulate global miRNA biogenesis in order to manipulate miRNA expression during infection (Bruscella et al. 2017). Thus, we sought to profile changes in the transcript levels of components of the miRNA biogenesis pathway, *dicer1* and *drosha*, in response to poly(I:C) or FV3. RT-qPCR analysis revealed downregulation of *dicer1* transcripts in response to poly(I:C) (24 h) as well as FV3 (24 and 72 h) (**Fig. 6a**). In addition, *drosha* transcripts were downregulated in response to FV3 infection at 72 h (**Fig. 6b**). Together, these results suggest that miRNA biogenesis is partially repressed in response to viral infection.

**Fig. 6.**
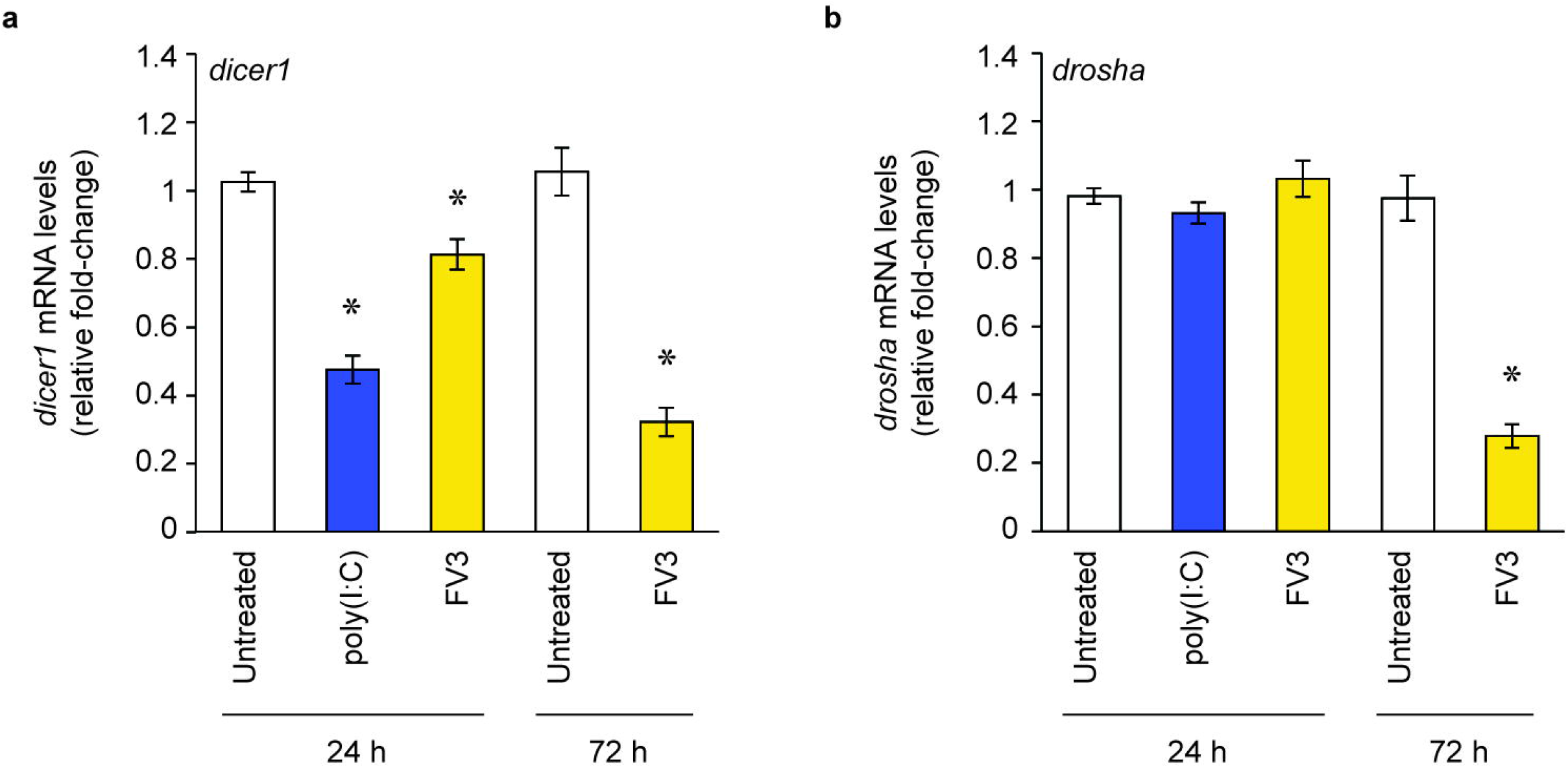
Fold-change in *dicer1* and *drosha* mRNA transcript levels in response to poly(I:C) treatment or FV3 infection of Xela DS2 cells. cDNA generated from total RNA isolated from untreated, poly(I:C)-treated, and FV3-infected Xela DS2 cells was subjected to RT-qPCR analysis targeting *dicer1* (**a**) and *drosha* (**b**) transcripts. RT-qPCR data were analyzed using the ΔΔCt method using xla-miR-16a-5p (levels unchanged by any treatment based on RNA-seq data) as an internal control. Data is depicted as mean ± standard error (*n* = 3), and values corresponding to treatment groups are presented relative to their respective untreated control, which was set to a value of 1. Statistical significance was assessed using a one-way ANOVA paired with Dunnett’s post-hoc test (24 h) or an unpaired Student’s *t*-tests (72 h). A *p* value < 0.05 was considered statistically significant and asterisks (*) denote statistical difference from the time-matched untreated control.

## Discussion

### Modeling of conserved and robust antiviral responses involving miRNAs

We uncovered 80 *X. laevis* miRNAs that are differentially expressed in response to poly(I:C) and/or FV3 in a frog skin epithelial-like cell line, suggesting for the first time that frog miRNAs are key regulators of antiviral immunity. We profiled changes in miRNA expression in response to poly(I:C) to model robust antiviral responses, as poly(I:C) induces potent antiviral responses and is an inert molecule (non-immunoevasive). Poly(I:C) treatment altered the levels of 65 *X. laevis* miRNAs, which may play a role in epigenetic regulation of host gene expression in response to exogenous dsRNA. Poly(I:C) treatment induced changes in miRNAs that are predicted to target important regulators of antiviral signaling pathways such as cGAS-STING (e.g. *stim1*), RIG-I/MDA5 (e.g. *trim25*), TLR signaling (e.g. *irak2, irf5, ripk1*, and *tab2*), type I IFN signaling (e.g. *ifnar1* and *stat1*), NF-κB-induced genes (e.g. *tp53*, and *sod2*) and ISGs (e.g. *adar* and *isg20l2*). However, it is important to note that currently available *X. laevis* genome annotations remain incomplete, and immune genes are particularly under-annotated. Thus, while target predictions were performed with the most up-to-date annotation, our target prediction analysis may represent an underestimation of immune gene targets. Additionally, we restricted our analyses to 3’ UTRs, as although miRNAs have recently been shown to interact with 5’ UTRs and cds, the 3’ UTR is the location most often targeted by miRNAs (Lee et al. 2009; Fang and Rajewsky 2011). Nevertheless, targeting of these antiviral pathways is reflective of what has been observed in other vertebrates, where miRNAs that are differently expressed during immune responses have been found to target orthologs of these same immune genes (Boosani and Agrawal 2016). For example, miR-146a is induced by vesicular stomatitis virus in murine peritoneal macrophages (Hou et al. 2009) and targets *irak2* (mice (Hou et al. 2009)), *irf5*, and *stat1* (humans (Tang et al. 2009)); miR-155 is induced during inflammatory responses in humans and mice (O’Connell et al. 2007; Tili et al. 2007) and targets *ripk1* (Tili et al. 2007) and *tab2* (Imaizumi et al. 2010); miR-208b and miR-499a-5p are induced in human hepatocytes in response to hepatitis C virus infection and target *ifnar1* (Jarret et al. 2016); and miR-30a is induced in HeLa cells in response to coxsackievirus B3 and targets *trim25* (Li et al. 2020). Our data sheds light on conserved epigenetic responses to viral dsRNA and the potential for involvement of miRNAs that target a core set of conserved antiviral genes. For example, it has been shown in mammals that poly(I:C) induces Sod2 expression to combat reactive oxygen species (ROS), which are associated with viral infection (Molteni et al. 2014). However, the mechanism through which poly(I:C) induces Sod2 remains unknown. We identified a miRNA (xla-miR-106-5p) that targets *sod2* and is downregulated in response to poly(I:C). Similarly, human hsa-miR-106a-5p is also predicted to target Sod2. Thus, repression of xla-miR-106-5p may be a conserved mechanism through which Sod2 is induced to combat ROS accumulation in response to viral infection. In line with this, oxidative stress has been shown to repress miR-106-5p in humans (Tai et al. 2020), and downregulation of miR-106-5p has separately been shown to restore Sod activity to inhibit apoptosis and oxidative stress in rats (Li et al. 2017). Together, these observations suggest that miRNA-mediated mechanisms of regulating innate immunity are highly conserved, even in distantly related vertebrates.

### Immunoevasive viruses produce host miRNA profiles distinct from those associated with robust and effective antiviral responses

FV3 is immunoevasive (Grayfer et al. 2015; Robert et al. 2017) and does not elicit robust antiviral responses in Xela DS2 skin epithelial-like cells (Bui-Marinos et al. submitted). Thus, we sought to profile changes in miRNA expression in response to FV3 for comparison to poly(I:C)-induced miRNA alterations to identify potential miRNA-based mechanisms of immunoevasion. Accordingly, FV3-infection is associated with changes in the expression of fewer miRNAs compared to poly(I:C), such that only 34 of the 65 miRNAs altered by poly(I:C) treatment were also affected by FV3 infection. Thus, the 31 miRNAs that are altered by poly(I:C) treatment and not FV3 infection, along with the 15 miRNAs that are altered by FV3 infection and not poly(I:C) treatment, may play a role in FV3’s ability to evade host immune systems in skin epithelial cells. In addition, some miRNA changes evident at 24 h post-poly(I:C) treatment were not evident until 72 h post-FV3 infection, and fewer of the differentially expressed miRNAs in FV3-infected Xela DS2 cells are predicted to target immune genes compared to those altered by poly(I:C). These findings suggest that FV3 infection may be associated with a dampened and/or delayed immune response, which is in accordance with the theory that FV3 possesses immunoevasion capabilities. To our knowledge, only one other study has simultaneously profiled changes in miRNA expression in response to poly(I:C) and an immunoevasive virus [porcine reproductive and respiratory syndrome virus (PRRSV)] (Wu et al. 2019). Similar to our observations, PRRSV infection of porcine alveolar macrophages resulted in alterations to only 33 of the 197 miRNAs found to be altered by poly(I:C) treatment (Wu et al. 2019). However, in contrast to the findings of this study, which did not detect any miRNAs altered by PRRSV infection that were not altered by poly(I:C) treatment, we detected differential expression of a handful of miRNAs in response to FV3 infection that were unaffected by poly(I:C) treatment. Of particular interest are two novel *X. laevis* miRNAs with no known orthologs in other species (novel-xla-miR-16-5p and novel-xla-miR-16-3p) that are strongly induced (150-fold and 342-fold, respectively, according to RT-qPCR) in response to FV3 but not poly(I:C). These novel miRNAs originate from the same pre-miRNA and are both predicted to target important genes involved in innate antiviral immunity, including *tgfbr2* (transforming growth factor beta receptor 2; novel-xla-miR-16-5p) and *trim14* (tripartite-containing motif 14; novel-xla-miR-16-3p). Tgfbr2 is a receptor for the well-studied TGF-β anti-inflammatory cytokine and viral induction of novel-xla-miR-16-5p may result in dampening of TGF-ß signaling, preventing the host from limiting inflammatory responses to protect against cellular damage caused by excessive inflammation. Trim14 indirectly promotes type I IFN production by stabilizing cGAS (Chen et al. 2016). In the absence of Trim14, type I IFN antiviral responses are impaired (Chen et al. 2016). Viral induction of novel-xla-miR-16-3p may dampen host innate antiviral responses by repressing type I IFN signaling. Thus, the possibility exists that these miRNAs are induced by FV3 infection to alter host immune function in skin epithelial cells.

To our knowledge, only one other study has profiled changes in host miRNAs in response to ranavirus infection. Meng et al. (2018) profiled *Andrias davidianus* (Chinese giant salamander) miRNA responses to Chinese giant salamander iridovirus infection of spleen tissue and discovered numerous differentially expressed miRNAs predicted to target genes involved in important antiviral signaling pathways such as TLR, RIG-I/MDA5, and JAK-STAT pathways (Meng et al. 2018). Despite the similarities to pathways targeted in our study, only two miRNA families found to be differentially expressed in the study by Meng et al. (2018) were also found to be differentially expressed in response to FV3 in our study (miR-200 and miR-203, both downregulated). Thus, while the antiviral pathways targeted by miRNAs in response to different ranaviruses may mirror one another, it appears as though regulation of these pathways may be achieved using different miRNAs. Nevertheless, it will be important for future studies to verify that the differentially expressed miRNAs identified in our study function in antiviral responses and impact viral replication.

In addition to targeting host antiviral genes, several differentially expressed miRNAs are predicted to target viral transcripts corresponding to a myristoylated membrane protein (FV3 ORF 53R), LCDV1 orf2-like protein, and hypothetical proteins. FV3 ORF 53R has been suggested to interact with the host cellular membrane and is required for virion assembly (Whitley et al. 2010), and thus may represent an ideal molecule for the host immune system to target. It is important to note that our target prediction analyses were performed with FV3 cds, as the current FV3 genome annotation lacks defined 3’ UTRs. While viral cds are frequently targeted by host miRNAs (Wang et al. 2012; Zheng et al. 2013; Ho et al. 2016), our FV3 target predictions may represent a fraction of the total FV3 mRNAs targeted by host miRNAs.

### Repression of miRNA biogenesis as a facet of both effective immune responses and immunoevasion

Recent studies have shown that transient inhibition of miRNA biogenesis is critical during the activation of IFN responses in HeLa cells, and leads to reduced levels of specific miRNAs, many of which target innate immune genes (Witteveldt et al. 2018). Accordingly, our study revealed that the majority of differentially expressed miRNAs that target immune genes are downregulated rather than upregulated, and both poly(I:C) treatment and FV3 infection repressed *dicer1* transcript levels. In line with this, studies have detected repression of *dicer1* but not *drosha* in response to viral infection of human, non-human primate, and murine cells (vaccinia virus infection of HeLa, Vero, and 3T3 cells) (Grinberg et al. 2012). Therefore, transient Dicer repression may constitute a conserved antiviral response. Interestingly, we observed stark downregulation of *drosha* transcript levels in response to FV3 but not poly(I:C), which raises the possibility that Drosha repression may be linked to immunoevasion tactics employed by FV3. Accordingly, studies have demonstrated that during infection of human 293T cells with Sindbis virus or vesicular stomatitis virus, Drosha is shuttled to the cytoplasm and repurposed to cleave viral RNA to inhibit viral replication (Shapiro et al. 2014). Thus, it stands to reason that immunoevasive viruses may be equipped to restrict Drosha function to facilitate viral replication.

### Expansion of the frog miRNAome

Our sequencing efforts expanded the number of currently annotated *X. laevis* miRNAs from 247 to 380. Of the 133 novel *X. laevis* miRNAs we identified in Xela DS2 cells, 108 have never been detected in any species before. While several novel miRNAs were differentially expressed in response to poly(I:C) or FV3, the majority were not. A large portion of the novel miRNAs that were not altered by poly(I:C) or FV3 originate from repetitive regions found across multiple chromosomes and may thus function in controlling repetitive elements. Indeed, miRNAs have been recently characterized as emerging regulators of transposable elements (Pedersen and Zisoulis 2016). Thus, our findings are likely to stimulate exciting studies in research areas outside of innate immunity.

While we significantly increased the number of annotated *X. laevis* miRNAs, it is important to note that our novel miRNA discovery efforts have likely not completed our map of the *Xenopus* miRNAome, as miRNAs are often exclusively expressed in specific cell types, tissue types, developmental stages, or cellular contexts (e.g. infection) (Bartel 2004). Additionally, our novel miRNA discovery pipeline involved removing reads with < 10 counts in a given library, which may have impaired our ability to discover novel miRNAs that are expressed at low levels. However, removing reads with counts < 10 improves the signal-to-noise ratio (Law et al. 2016) which enables the discovery of high confidence miRNAs (Friedländer et al. 2012), and we felt this benefit outweighed the limitation of missing some novel miRNAs.

### Concluding remarks

To our knowledge, this study is the first to highlight the role of miRNAs in antiviral defenses within frog skin epithelial cells and sheds light on evolutionarily conserved epigenetic mechanisms of regulating antiviral responses in vertebrates. We identified “typical” antiviral responses involving miRNAs in frogs and uncovered differences between these typical responses and miRNA-mediated responses to the immunoevasive FV3. Our findings further our understanding of frog-FV3 interactions by identifying miRNAs as an added layer of antiviral gene regulation in frogs, and uncovering the possibility that ranaviruses may hijack specific host miRNAs to facilitate immunoevasion. Future functional studies will be instrumental in solidifying these initial observations, and similar studies in other cell types and tissues will be crucial to uncovering cell type- and tissue-specific antiviral miRNA responses in frogs.

## Supplementary material

**Fig. S1**. Characteristics of samples used for small RNA-seq. (**a**) Representative phase-contrast images of Xela DS2 cells from which RNA was isolated for small RNA-seq. Scale bar = 100 μm, *n* = 3 independent trials. (**b**) Viral titres (Log_10_ TCID_50_/mL) associated with FV3-infected Xela DS2 cell supernatants at 24 h and 72 h (*n* = 2). (**c**) RT-PCR detection of the cel-miR-39 spike-in miRNA as a means of confirming of the ability to detect miRNAs in each RNA sample to be sequenced. Detection of *actb* mRNAs is presented as an internal control. (**d**) Bioanalyzer analysis of RNA quality prior to sequencing. n.d. = not detected, kb = kilobase, RIN = RNA integrity number.

**Table S1**. RT-PCR and RT-qPCR primers used in this study

**Table S2**. Novel *X. laevis* miRNAs

**Table S3**. Predicted FV3 targets of known and novel *X. laevis* miRNAs

**Table S4**. Predicted endogenous targets of *X. laevis* miRNAs

