## Supplementary figures and images for "Epigenetic regulation of frog innate immunity: Discovery of frog microRNAs associated with antiviral responses and ranavirus infection using a *Xenopus laevis* skin epithelial-like cell line"

### Supplemental Figure 1

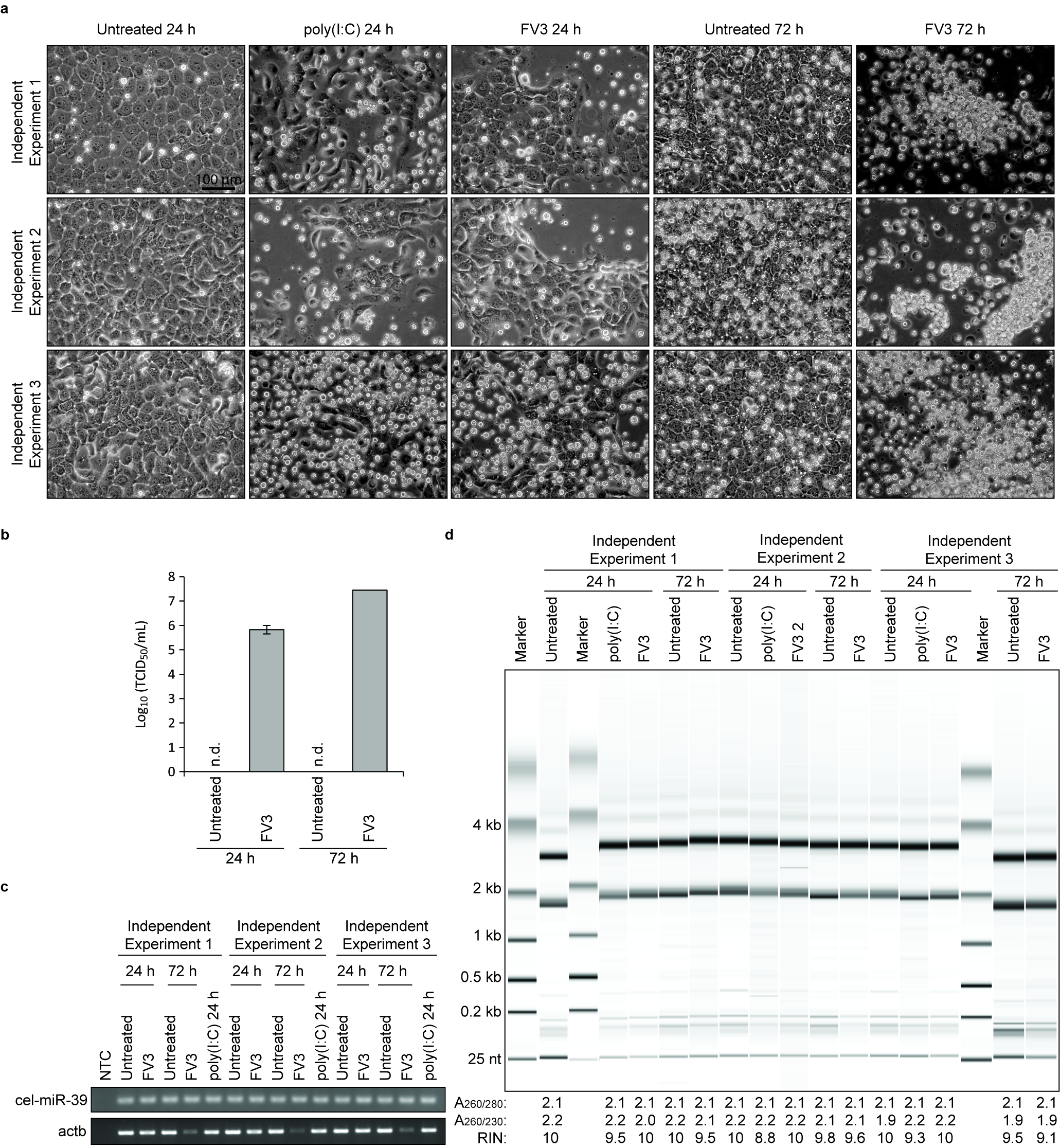
